# Protein aggregation capture on microparticles enables multi-purpose proteomics sample preparation

**DOI:** 10.1101/447904

**Authors:** Tanveer S. Batth, Maxim A.X. Tollenaere, Patrick L. Rüther, Alba Gonzalez-Franquesa, Bhargav S. Prabhakar, Simon Bekker-Jensen, Atul S. Deshmukh, Jesper V. Olsen

## Abstract

Universal proteomics sample preparation is challenging due to the high heterogeneity of biological samples. Here we describe a novel mechanism that exploits the inherent instability of denatured proteins for non-specific immobilization on microparticles by protein aggregation capture. To demonstrate the general applicability of this mechanism, we analyzed phosphoproteomes, tissue proteomes, and interaction proteomes as well as dilute secretomes. The findings presents a practical, sensitive and cost-effective proteomics sample preparation method.

## MAIN

Dedicated sample preparation for shotgun proteomics is essential for removing impurities and interfering species which may affect peptide chromatography, ionization during the electrospray process, and sequencing by mass spectrometers. To represent the in-vivo state of the global proteome including membrane-bound proteins, it is of high importance to ensure complete lysis of cells and tissues prior to protease digestion. This typically requires strong detergents that are difficult to remove afterwards, however crucial in order to avoid signal interference during MS analysis. Considerable developments have been made based on a variety of different biochemical principles which utilize filters, traps, or protein precipitation techniques which address different sample types^1,2,3^. However, a primary challenge remaining is the development of a universal sample preparation method that has the potential to scale across different sample amounts, which typically range from ng to mg of starting material. Moreover, such a method needs to be compatible with different lysis buffers, biological material (i.e. cell lines, tissues), robust, reproducible, cost effective, and perhaps above all; practical. Although several methods have been developed to individually address different proteomics sample preparation challenges, a simple solution spanning all sample types remains elusive. Here we report a mechanism, termed protein aggregation capture (PAC), which utilizes the phenomenon of non-specifically immobilizing precipitated and aggregated proteins on any type of sub-micron particles irrespective of their surface chemistry. We explore the fundamental process underlying this phenomenon and determine the optimal parameters leading to effective sample preparation for shotgun proteomics analysis by mass spectrometry of different sample types. Our developments demonstrate the potential for low cost, simple, robust and sensitive sample preparation procedures for proteomics analysis, which can be easily implemented in any setting with great potential for full automation.

Our hypothesis was based on a series of reports and observations that lead us to conclude the mechanism of non-specific aggregation, initially on magnetic beads with carboxyl group surface chemistries. Carboxyl coated magnetic beads have been reported for sensitive proteomics sample preparation as an alternative to other approaches such as FASP with limited starting material^4^. The binding mechanism was attributed to hydrophilic interactions (HILIC)^5^ with the carboxyl surface groups and the method was termed “SP3”. The method initially suffered from low recovery and reproducibility, and thus was not widely adopted although recent improvements of the protocol, such as pH control, rendered it more practical^6^. As HILIC principles dictate preferential polar and ionic interactions under non-aqueous conditions, protein interaction to the carboxyl surface of the beads was hypothesized to be induced by the addition of acetonitrile to the protein lysate. However, we observed that stringent binding of proteins to the microspheres could not be reversed under aqueous conditions even with extended washing, which would contradict HILIC principles (Figure 1A, supplementary Figure 1A). Proteins however could be released in solubilization buffers such as lithium dodecyl sulfate (Figure 1A). We therefore wondered whether protein immobilization on magnetic microspheres was driven by aggregation of insoluble proteins on microspheres rather than hydrophilic interactions. To test this, we treated native protein lysates either by incubation at room temperature (25°C) where proteins should stay in their native state, or induced aggregation by three different known mechanisms, these being organic solvent (acetonitrile; 70% final), high temperature (80°C for 5 minutes), or high salt (2.5 M ammonium sulfate), followed by the addition of magnetic carboxyl microspheres. Immobilization of aggregated and insoluble proteins was only observed under the three conditions known to induce aggregation indicating that protein aggregation was essential to the underlying mechanism of protein capture (Figure 1B). Importantly, the induced protein aggregation was very effective - especially using acetonitrile - as judged by the little protein amounts remaining in the supernatants. We subsequently investigated the role of microsphere surface chemistry on protein immobilization and found no impact on protein aggregation irrespective of microsphere surface chemistry including those containing hydrophobic C_18_ surfaces, which contradicts the HILIC driven hypothesis of on-bead protein immobilization (Figure 1C). To rule out the role of the magnetic properties of the microspheres leading to immobilization, we tested protein aggregation on porous 3μm C_18_ hydrophobic beads, which are typically utilized for packing reversed-phase nano-columns and found similar immobilization mechanisms (Supplementary Figure 1B). We further examined whether coated smooth surface microspheres were essential for protein immobilization by inducing protein aggregation (with acetonitrile) on fine iron powder microparticles (grain size 5-9μm) and observed aggregation in a similar manner (Supplementary Figure 1C). We next inquired whether protein aggregation on microspheres was a function of microparticle surface area by gauging protein aggregation at very low microsphere concentrations relative to a constant concentration of protein lysate at 0.25 μg/ul (Figure 1D). Although very low amounts of beads were sufficient to aggregate proteins from solution (Figure 1D), we found the structural integrity of visible protein-bead precipitate to be unstable when the bead to protein ratio was less than 1:4, leading to dispersion of small aggregated pieces in solution upon mild disruption. Conversely, solutions with low protein concentrations (<0.075 μg/μl) were found to require higher bead to protein ratios for efficient capture and recovery (Supplementary Figure 1D, E). These results indicate that solutions with higher protein concentrations can aggregate on minute amounts of microparticles, however lower protein concentration require relatively higher amounts of microparticles in order to effectively capture aggregated proteins. Collectively the data suggest that microparticle surface (irrespective of surface chemistry) acts as a nucleation site or carrier to induce a immobilization cascade of insoluble protein aggregates, which ultimately serve to tightly maintain a microparticle-protein structure (Figure 1E).

**Figure 1.**
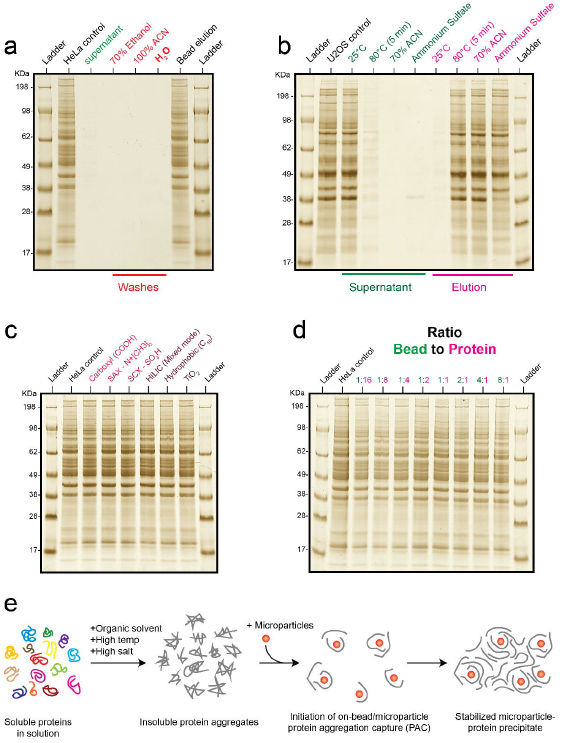
Elucidating the mechanism of protein aggregation capture on microparticles. A) The hypothesis of HILIC based bead interactions was tested by inducing bead – protein interaction on 20μg of HeLa protein lysate (in 1% SDS) with the addition of acetonitrile (70% final concentration) and microparticles (20μg) separated by magnet. The resulting supernatant was analyzed by SDS gel electrophoresis. Beads were sequentially washed three times with the indicated buffers. All washes were analyzed by SDS-PAGE for protein elution by beads including washes by milli-Q water and the lanes indicated in red. After bead washing, LDS buffer was added to the beads (see materials section) and analyzed. B) U2OS protein lysates (in 0.1% NP-40) were treated with different conditions as indicated in green. Carboxyl coated magnetic beads were added to the lysates after treatment and the resulting supernatant analyzed by SDS-PAGE. LDS buffer was similarly added to the beads and the resulting supernatant analyzed by SDS-PAGE. C) Acetonitrile was added to HeLa lysate (in 1% SDS) to a final concentration of 70% and equal amounts of microparticles with different surface chemistries were added to the lysates and the supernatant removed. LDS buffer was added to the microparticles and the resulting supernatant analyzed by SDS-gel after removal by magnet. D) Aggregation of equal amount of HeLa lysate (20μg at 0.25μg/μl after addition of acetonitrile) was induced in a similar fashion as indicated above and carboxyl coated microparticles were added to the lysate at different amounts as indicated in the figure. The supernatant was removed and the LDS buffer was added to the different samples and analyzed by SDS-PAGE after separation of microparticles by magnet. E) The hypothesized model for protein aggregation capture (PAC) on microparticles is illustrated based on the above observations.

We assessed the impact on trypsin digestion efficiency of immobilized proteins on microparticles and compared it to a commonly-used chaotropic agent based in-solution digestion protocol^7^. Analyzing all samples by single shot nanoflow liquid chromatography tandem mass spectrometry (LC-MS/MS) we found significantly reduced number of missed tryptic cleavages between the two methods at different Lys-C and trypsin ratios (Figure 2A, supplementary figure 2A). These findings imply the possibility to considerably reduce proteomics sample preparation costs as proteases typically constitute one of the largest expense of the workflow prior to MS analysis. As previously determined by missed cleavages across different digestion protocols^8^, our results indicate that 10-20x reduction in trypsin and Lys-C usage (1:500 - 1:1000 ratio) leads to comparable missed cleavage rates (below 30%) to the standard 1:50 trypsin-to-protein ratio utilized in most proteomics studies (Figure 2A, supplementary figure 2A).

**Figure 2.**
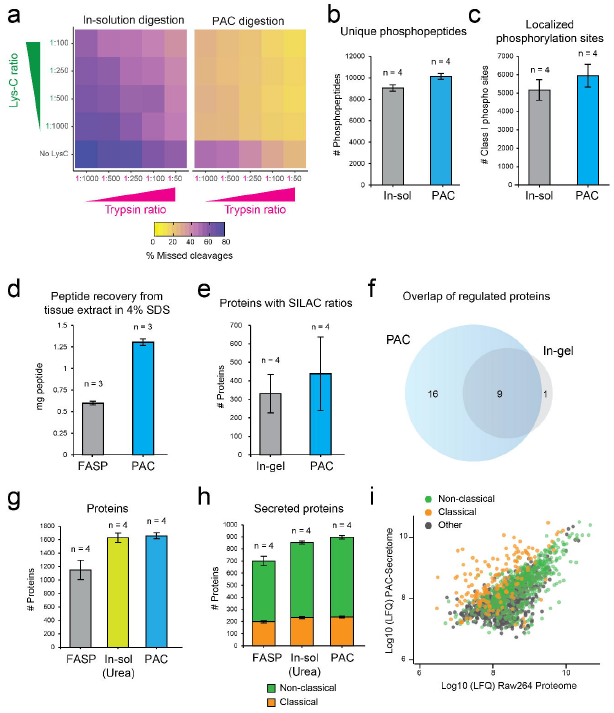
Exploring the boundaries of PAC for proteomics analysis of different sample types. A) Percentage of peptides containing missed cleavages at arginine or lysine after digestion with trypsin and lys-c proteases in different combinations and ratios are displayed by heatmap. Missed cleavage rates were investigated on lysates prepared by protein aggregation on microspheres or in-solution digestion. B) Average number of unique phosphopeptide variants were counted (after removal of contaminant or reverse hits as defined by MaxQuant analysis) for the different experiments. Phosphopeptides were tallied after enrichment from lysates prepared with in-solution or PAC digestion. C) Average number of phosphorylation sites with high localization probabilities (as defined by site localization probability ≥0.75) are presented between the two different methods. D) Average peptide recovery after protease digestion of equal amounts (∼1.8 mg) of mouse skeletal tissue prepared using the FASP or PAC protocol measured by nanodrop absorbance at 280/260nm. E) The average number of proteins with SILAC ratios were counted between the two different experiments after removal of contaminating proteins and reveres hits. F) Overlap of statistically regulated protein between two preparation methods as determined by t-test with permutation-based FDR of 0.05 with 250 randomizations and s0 of 0.1. G) Number of proteins containing LFQ intensities from the different secretome workflows. H) Number of secreted proteins between the different experiments (see Methods). I) Median Log10 transformed LFQ intensities of the proteins identified in PAC are plotted against those identified in the base proteome of Raw264.7 cells. Classical and non-classical secreted proteins are highlighted. *Error bars represent standard deviation in all cases.

Post-translational modifications (PTMs) such as site-specific phosphorylation can rapidly modulate the function of proteins by changing their enzymatic activity, subcellular localization, turnover, and interaction partners^9^. It is therefore important to develop proteomics methods that enable global analysis of phosphoproteomes in a robust, reproducible and sensitive manner. We examined whether the analysis of protein phosphorylation status is affected by the on-bead protein aggregation capture workflow on serum stimulated HeLa cells. Phosphopeptides were enriched by magnetic Ti-IMAC beads and the eluates analyzed by LC-MS/MS in turn. The results demonstrate no impact in the number of identified phosphorylated peptide variants and phosphorylation sites (Figure 2B, C). Importantly, >1000 more phosphorylated peptides and 779 localized sites were identified on average using the microsphere based method compared to standard in-solution digestion (Figure 2B, C). Moreover, high degree of overlap for localized sites was found between the two methods (Supplementary figure 2B) and no bias in the phosphopeptide enrichment was observed between the two experiments as we achieved an enrichment efficiency >99% for all replicates (Supplementary figure 2C).

We next evaluated the potential of utilizing magnetic microparticles for proteomics analysis of organs and tissues. This can be particularly challenging as it often requires harsh solubilization buffers for efficient protein extraction from hard and soft tissues. To test aggregation on microparticles we utilized skeletal muscle tissue samples from *mus musculus*. After homogenization and solubilization in 4% SDS lysis buffer, we examined protein recovery and digestion by using either the established filter-aided sample preparation (FASP) protocol or the PAC workflow using microspheres with sulfonic acid surface chemistry. Significantly higher peptide recovery (>2 fold) after Lys-C/trypsin digestion was observed using microspheres from initial starting material of 1.8 mg (as determined by tryptophan assay^10^) per replicate (Figure 2D). Incidentally, sample preparation with magnetic microspheres resulted in a cleaner peptide mixture as no polymer peaks were observed in the mass spectrometry analysis, leading to higher number of identified proteins and unique peptides in a single shot LC-MS/MS analysis (Supplementary Figure 2D, E).

Proteins rarely operate alone in the cell and their function is usually dependent on the protein complexes they are part of^11^. One potential application of magnetic microparticles is for the rapid analysis of protein-protein interactions (PPIs). Antibody-based pulldown in combination with mass spectrometry is a popular approach for elucidating protein-protein interactions (PPIs). We inquired whether a simplified microparticle based method for analyzing PPIs provides depth of coverage comparable to standard in-gel protocols. To test this we utilized a SILAC^12^ based setup for determining interactors of ZFP36 (Tristetraprolin or TTP), an RNA binding protein (Supplementary Figure 3A). On average we identified 100 more proteins (438) in a PAC based single shot MS analysis compared to a conventional in-gel digestion workflow (330) with 5 fractions per replicate (Figure 2E, supplementary Figure 3B). Crucially, we found good overlap between the two groups for IL-1b regulated TTP-interactors (Figure 2F). We identified known interactors of TTP upon IL-1b stimulation such as 14-3-3 subunits and RNA-binding factors which regulate stability such as UPF1 and PABC1/4^13^ (Supplementary figure 3C), as well as potential novel interactors which displayed interesting interaction dynamics with TTP upon IL-1b stimulation (Supplementary figure 3D).

MS-based analysis of cellular secretome holds enormous promise for the investigation of cellular communication. However, the analysis of cellular secretome presents several challenges as proteins are usually secreted in low concentrations making their detection in culture media (which are rich in salts and other compounds) difficult^14^. We sought to determine the applicability of microparticles for enriching secreted proteins in the background of cell culture media contaminants. To this end we benchmarked urea (in-solution) and filter (FASP) based methods as reported previously^15,16^, against PAC on microparticles for secreted proteins. As described above (Supplementary figure 2C, D), high concentration of microspheres (>300 μg/ml) were required in order to provide sufficient surface area for immobilization of dilute aggregated proteins. Protein aggregation on microspheres consistently identified the largest number of proteins and unique peptides resulting in the highest sequence coverage (Figure 2G, supplementary figure 4A, B). Although the low peptide recovery with FASP protocol led to the fewest number of protein identifications, the microsphere and FASP methods produced the cleanest peptide sample as determined by spectroscopy analysis (Supplementary Figure 4C, D). Using previously described computational workflow to predict potentially secreted proteins^16^, we found majority (>50%) of the detected proteins have been previously characterized as secreted proteins through the classical secretory pathway (via signal peptide) or other non-conventional pathways (Figure 2G)^17^. However, proteins secreted through the classical pathway were found at higher abundance in the media (Figure 2I). This includes low abundant cytokines such as Csf3 and Cxcl10 which were only identified using the PAC method, demonstrating sensitivity of the protocol. Thus, microparticle aggregation can enable simultaneous quantification of 100’s of secreted proteins in the background of complex cell culture media.

We have described the protein aggregation capture mechanism behind microsphere based protein immobilization. Understanding this in great detail have led to optimized protocols that outperform competing methods for proteomics sample preparation which include tissue proteomes, enriched subproteomes such as phosphoproteomes, low abundant immunoprecipitated and secreted proteins. We demonstrate that the developed PAC mechanism is extremely scalable and has further advantages such as very low cost, simplicity, and time effective. Limitations of the method could potentially include samples containing certain components (such as acids) which could prevent efficient aggregation on microparticle surfaces. Other large biomolecules such as DNA/RNA can also co-precipitate with proteins if not adequately removed. We hope awareness of this mechanism will lead to further novel developments and applications. Future developments could utilize microparticle surface functional group specificities for peptide level enrichment/fractionation following non-specific aggregation at the protein level.

## METHODS

### Reagents

Chemicals were purchased from Sigma-Aldrich unless otherwise specified. Sera-mag carboxyl magnetic beads (cas # 45152105050250 and cas # 65152105050250) were purchased from GE-Healthcare. SIMAG-Sulfon (cas # 1202), SiMAG-Q (cas # 1206), and SiMAG-Octadecyl (cas # 1301) magnetic beads were all purchased from Chemicell GmbH. Magnetic HILIC, TiO_2,_ and Ti-IMAC magnetic beads were purchased from ReSyn Biosciences. Carbonyl-iron powder was purchased from Sigma-Aldrich (cas # 44890).

### Cell culture

Human bone osteosarcoma epithelial (U2OS) and human epithelial cervix carcinoma (HeLa) adherent cells were grown in DMEM media (Gibco) supplemented with fetal bovine serum (Gibco) at 10% final. The media also contained penicillin (Invitrogen) at 50 U/mL and streptomycin (Invitrogen) at 100 μg/mL. Cells were grown in a humidified incubator at 37°C with 5% CO_2_. In all cases, cells were grown to 80-90% confluency before harvesting with different lysis buffers in Nunc petridishes (100 or 150mm diameter).

To generate stably expressing GFP-TTP cells under a doxycycline inducible promoter, ZFP/TTP was gateway cloned into a pCDNA4/TO/GFP expression vector by gateway cloning (Thermo Fisher), and co-transfected with pcDNA6/TR (Thermo Fisher) into U2OS cells. Cells were selected with zeocin and blasticidin for 14 days, after which individual clones were picked and screened for GFP-TTP expression. For SILAC labelling, cells were cultured in media containing either L-arginine and L-lysine (Light), L-arginine [^13^C6] and L-lysine [^2^H4] (Medium) or L-arginine [^13^C6-^15^N4] and L-lysine [^13^C6-^15^N2] (Heavy; Cambridge Isotope Laboratories).

RAW264.7 macrophage cells were derived from *mus musculus* and grown in 10% in DMEM media with 10% FBS in 150mm diameter Nunc petridishes. The media was removed and cells were washed with PBS prior to addition of phenol-red free DMEM media without serum, penicillin, and streptomycin. Cells were stimulated with lipopolysaccharids (LPS) with 1μg/ml for 4 hours. 400 μl of the media was removed and processed for secretome analysis and filtered through 0.22uM filter (Sartorius #16532) prior to further processing.

### Cell lysis and sample preparation

Cells lysis as presented in this study was performed with either one of the three buffers: 1) 6M guanidine hydrochloride in 100 mM Tris Hydrochloride (Life technologies) at pH 8.5, 2) 1% SDS in 100 mM 100 mM Tris Hydrochloride (pH 8.5) or 3) 0.1% NP-40 in 1X phosphate buffered saline solution (pH 7.4) containing β-glycerol phosphate (50 mM), sodium orthovanadate (10 mM), and protease inhibitor cocktail (Roche). In all cases, supernatant from adherent cell plates was removed and the cells were rinsed with ice cold 1X PBS prior to the addition of the lysis buffer.

Guanidine hydrochloride buffer was pre-heated to 99°C prior to the addition to the cell plates. After the addition of guanidine or SDS buffer, cells were manually collected and heated at 99°C for 10 minutes followed by sonication using a probe to shear RNA and DNA. For cells lysed using 0.1% NP-40 buffer, 1μl of benzonase (≥250U/ul) was added to the lysis solution for 1 hour on ice. The lysis solution was centrifuged at 5000 x G for 10 minutes and the supernatant was transferred to a new tube.

GFP-TTP immunoprecipitations were performed using GFP-Trap magnetic agarose beads (Chromotec) according to manufacturer’s instructions. Cell lysis and immunoprecipitations were carried out using low salt EBC lysis buffer (150 mM NaCl; 50 mM TRIS pH 7.5; 1 mM EDTA; 0,5% NP40).

### On-bead protein aggregation

Aggregation was induced by the addition of acetonitrile (unless stated otherwise) and magnetic microparticles were added to solution followed by brief mixing with pipette tip. The solution was allowed to settle for 10 minutes and beads were separated using a magnet for 60 seconds. Magnetic microspheres were retained by magnet and the supernatant was removed by vacuum suction. In the case of analysis by protein gel electrophoresis, the supernatant was transferred to new tubes. Beads were washed using acetonitrile or ethanol once followed by one wash with 70% ethanol. Organic solvent from bead washing was evaporated using speedvac. Samples and washes were prepared for analysis by protein gel electrophoresis (SDS-PAGE) by the addition of 4x LDS sample buffer (Thermo Fisher Scientific) to 1x final, and DTT (100 mM). Samples were heated for 10 minutes at 80°C. For eluting bead bound protein aggregates, LDS buffer (containing DTT) was added to magnetically separated beads and the mixture was heated for 10 minutes at 80°C. Heated beads in LDS buffer were separated by magnet and the supernatant was analyzed by SDS-PAGE or transferred to a new tube and stored at −20°C until SDS-PAGE analysis. Samples were loaded on NuPAGE 4-12% Bis-Tris protein gel (Thermo Fisher Scientific) and ran with 200 volts for 40 minutes. Gels were stained for 15 minutes using instant Blue (Expedeon) and destained overnight with Milli-Q water and later scanned on EPSON V750 PRO.

One limit of aggregating proteins on microparticles that we observed was low peptide recovery (after protease digestion) from protein lysates containing high concentration chaotropic salts such as guanidine hydrochloride (6M). We attributed this to phase separation upon addition of organic solvents such as acetonitrile in the aqueous sample buffer. This reduced protein aggregation as proteins were remained soluble in the aqueous phase containing guanidine hydrochloride. However, diluting the concentration of the salt containing buffer with water prior to organic solvent addition ameliorated phase separation as determined by peptide recovery after on-bead trypsin digestion (Supplementary Figure 4E). Alternatively, organic solvents with higher water solubility could be also be utilized to prevent phase separation.

### Lys-C and Trypsin digestion and peptide cleanup

Proteins were aggregated on microspheres and washed as described above. For on-bead digestion, 50 mM HEPES buffer (pH 8.5) was added to submerge microspheres. Proteins were reduced and alkylated with the 5mM tris(2-carboxyethyl)phosphine (TCEP) and 5.5mM 2-chloroacetamide (CAA) for 30 minutes. Lys-c (Wako chemicals) was added at ratio of 1:200 (to protein) and allowed to react for 1 hour at 37°C followed by the addition of trypsin at a ratio of 1:100 (unless specified otherwise). Trypsin digestion was allowed to occur overnight at 37°C. Beads were separated by magnet and the supernatant was transferred to new tube and acidified.

In-solution digestion with guanidine hydrochloride buffer was carried out under similar reduction and alkylation conditions. Lys-c was added to solution and allowed to react for 1 hour at 37°C. The concentration of guanidine hydrochloride concentration was reduced to >1M prior to the addition of trypsin for overnight digestion. Solution was acidified by with 1% trifluoroacetic acid (TFA) and centrifuged for 5 minutes at 5000 x G and the supernatant transferred to new tubes.

Peptide mixtures were clarified with solid phase extraction. Briefly, hydrophobic C18 sep-pak (Waters Corporation) were prepared by washing with acetonitrile and 0.1% TFA, followed by loading of the acidified peptide mixtures by gravity. Sep-paks were washed with 0.1% TFA and peptides were eluted using 50% Acetonitrile (0.05% TFA). Organic solvent was evaporated and peptides concentrated using a speedvac prior to MS analysis.

### Protein extraction from mouse skeletal tissue

We used skeletal muscle which were isolated for previously published study (Schonke et al 20018, Proteomics). Frozen gastrocnemius muscles were crushed using mortar and pestle. Powdered muscle was homogenized using Ultra Turrax T8 homogenizer (IKA Labotechnik) in 4% SDS buffer (100 mM Tris-HCl, pH 7.4). Protein lysates were boiled at 100 °C for 5 mins. Lysates were sonicated using a tip and centrifuged at 16000g for 10 mins, the supernatant was then frozen until further analysis.

### Filter-aided sample preparation (FASP)

Urea powder was added to 400uL of filtered cell culture supernatant for a final 2M concentration, and pH for digestion adjusted with 40uL Tris 1M pH 8.5. FASP protocol was adapted from as previously described^1^. Samples were heated for 10 min at 56°C and centrifuged (7000g, 10min). Following centrifugation steps were performed applying the same conditions. Ultracel-30 membrane filters (Millipore #MRCF0R030) were cleaned with 10% acetonitrile and 15% methanol, filters were centrifuged, and equilibrated with 200uL urea buffer (2M, 0.1M Tris, pH8.5), and centrifuged again. Samples were added into the filters and the filters were centrifuged and washed two times with urea buffer. Reduction was performed by 1uL of 0.5M TCEP in 100uL urea buffer. The device was centrifuged and alkylation was performed by 1uL of 550mM CAA in 100uL urea buffer for 30 minutes in the dark. Filters were centrifuged and 200uL urea buffer was added prior to another centrifugation. Subsequently, 4uL of 0.5 ug/uL lys-c in 40uL urea buffer was added for 3h at 37°C with gentle orbital shaking. 4uL of 0.5ug/uL trypsin was added for an overnight digestion in the wet chamber at 37°C with gentle orbital shaking. 1.5mL eppendorf tubes were cleaned with absolute methanol and air dried, prior to inserting the filter device, which was then centrifuged. Subsequently 40uL of milli-Q water was added followed by centrifugation. The enzymatic digestion was stopped by acidifying the sample to pH<2.5 with TFA. StageTipping was performed right after.

### In-solution urea digestion sample preparation

All following chemicals have the same references and concentrations as in the FASP sample preparation. Urea powder was added to 400uL of filtered (0.22uM) cell culture supernatant for a final 2M concentration, and pH for digestion adjusted with 40uL Tris 1M pH8.5. 100% v/v. TCEP was added and the tubes were incubated for 30 minutes. Subsequently, samples were incubated with CAA for 20 minutes in the dark. The digestion step included addition of lys-c, incubation during 3h, followed by the addition trypsin (0.5ug/uL), and incubation overnight at room temperature. The enzymatic digestion was stopped by acidifying the sample to pH<2.5 with TFA. Samples were desalted and concentrated using Stage-Tips.

### In-gel digestion

In-gel protein digestion and downstream processing was performed as described earlier^18^.

### Enrichment of phosphorylated peptides

Adherent HeLa cells were grown as described above. Cells were washed and serum starved (DMEM without FBS) for 4 hours followed by 10 minute stimulation with FBS (10%). Cells were rapidly washed and lysed using guanidine hydrochloride buffer as described above. Protein concentration was estimated using tryptophan assay. Lys-C and trypsin digestion was carried out as described above. Peptides were clarified using SPE as described above with the exception that the peptide mixture was not concentrated using a speedvac. Small aliquat representing 5% was removed for determining peptide concentration using nanodrop which was estimated to roughly 200μg.

Ultra high phosphopeptide enrichment efficiency was achieved using Ti-IMAC magnetic beads (ReSyn Biosciences) with slight modification to the manufacturer protocol. Beads were prepared according to manufacturer instructions and loading buffer (80% acetonitrile, 1M glycolic acid, 5% TFA) was added to eluted peptide mixture in Protein LoBind 96 well plate (Eppendorf) at 1:1 volume ratio. 200μg of Ti-IMAC beads (10μl) were added to the peptide mixture and the binding occurred at 1400 RPM on a benchtop 96 well plate shaker at room temperature for 20 minutes. Magnetic beads were separated from the supernatant with a 96 well plate magnet stand (Ambion) and the supernatant was removed using vacuum suction via gel loader tip. Beads were washed with 400 μl of loading buffer for 1 minute and the supernatant was removed in a similar fashion. Beads were washed twice for 2 minutes with 400 μl of washing buffer (80% acetonitrile, 1% TFA) for 2 minutes each. Beads were washed twice for 2 minutes with 400 μl of washing buffer 2 (10% acetonitrile, 0.2% TFA). Following removal of supernatant from the last wash, 80 μl 1% ammonium hydroxide was added to the beads for 20 minutes at 1400 RPM. The supernatant was transferred to Protein LoBind tubes (Eppendorf) and the step was repeated twice, each time with 80 μl of 1% ammonium hydroxide for a final volume of 240 μl. 60 μl of 10% formic acid was added to the final solution and the sample was speedvac’d for 30 minutes at 60°C. Phosphopeptide containing solution loaded onto C18 STAGE-tips where the phosphopeptides were loaded and washed. The STAGE-tips were stored at 4°C until elution and analysis by MS.

### Mass spectrometry and liquid chromatography

Samples were injected on a 15 cm nanocolumn (75μM inner diameter) packed with 1.9μM C_18_ beads (Dr. Maisch GmbH) using an Easy-LC 1200 (Thermo Fisher Scientific). Peptides were separated and eluted from the column with an increasing gradient of buffer B (80% acetonitrile, 0.1% formic acid) at a flow rate of 250 nL/minute.

All samples were analyzed on a Q-Exactive HF-X (Thermo Fisher Scientific) mass spectrometer coupled to EASY-nLC 1200. With the exception of two replicates of in-gel TTP pulldown and one replicate from PAC TTP pulldown experiment, which were analyzed on a Lumos (Thermo Fisher Scientific) mass spectrometer with similar scan settings. The mass spectrometer was operated in positive mode with TopN method.

### Data analysis

Raw files generated from LC/MS/MS experiments were analyzed using MaxQuant (1.6.1.1) software^19^. Samples generated from human cell lines (HeLa and U2OS) were searched against the reviewed Swiss-Prot human proteome (proteome ID: UP000005640, release date March 2018). Samples generated from mouse cell lines (Raw264.7) and tissue were searched against the *mus musculus* reviewed Swiss-Prot proteome (proteome ID: UP000000589, release date October 2018). All searches were performed with carbamidomethyl of cysteines as a fixed modification while methionine oxidation and protein n-terminus acetylation were set as variable. Phosphorylation of serine, threonine, and tyrosine were set as variable modification for analysis of phosphopeptide enriched samples using Ti-IMAC. A false discovery rate (FDR) of 1% was utilized for peptide spectral matches, peptides, and proteins. Match between runs feature was utilized only for the analysis of phosphopeptide enriched samples.

Secretome analysis was performed using Perseus 1.5.2.6^20^. Proteins were annotated using UniprotKB keywords and gene ontology cellular components (GOCC). Proteins with keyword “Signal” were defined as classically secreted. Proteins annotated with Uniprot keyword “Secreted” and GOCC terms “extracellular region”, “extracellular space”, or “extracellular region part” were defined as non-classically secreted.

## ACKNOWLEDGEMENTS

We would like to thank Prof. Christian Kelstrup for constructive discussion and feedback. Work at The Novo Nordisk Foundation Center for Protein Research (CPR) is funded in part by a generous donation from the Novo Nordisk Foundation (Grant number NNF14CC0001). The proteomics technology developments applied was part of a project that has received funding from the European Union’s Horizon 2020 research and innovation programme under grant agreement No 686547 (MSmed). We would like to thank the PRO-MS Danish National Mass Spectrometry Platform for Functional Proteomics and the CPR Mass Spectrometry Platform for instrument support and assistance. T.S.B is funded by the HOPE project grant from the Novo Nordisk Foundation (Grant number NNF17SA0027704). P.L.R. is supported by the Marie Skłodowska-Curie European Training Network (ETN) “TEMPERA”, a project funded by the European Union’s EU Framework Programme for Research and Innovation Horizon 2020 (Grant Agreement number 722606). Work in the Bekker-Jensen lab was supported by grants from the Lundbeck Foundation, The NEYE Foundation and The Danish Medical Research Council.

### AUTHOR CONTRIBUTIONS

T.S.B and J.V.O conceived and designed the project. T.S.B developed the workflows and carried out the experiments. M.A.X.T generated the stable U2OS cell lines expressing GFP-TTP and performed the experiments and GFP-TTP pulldowns. M.A.X.T and S.B-J provided the Raw264.7 cell lines used for secretomics analysis and assisted in the data analysis of SILAC GFP-TTP pulldown experiments. P.L.R carried out the experiment for determining protease digestion efficiency and performed the data analysis. A.G.F and A.S.D performed the secretome experiments using FASP and in-solution workflows and assisted in the downstream data analysis of secreted proteins. A.S.D provided the *mus musculus* skeletal muscle tissue and assisted with the experiments. B.S.P performed the SDS-PAGE experiments. T.S.B wrote the initial draft of the manuscript and all authors contributed to the manuscript.

### CONFLICTS OF INTEREST

Authors declare no competing interests.

---

**Supplementary Figure 1.**
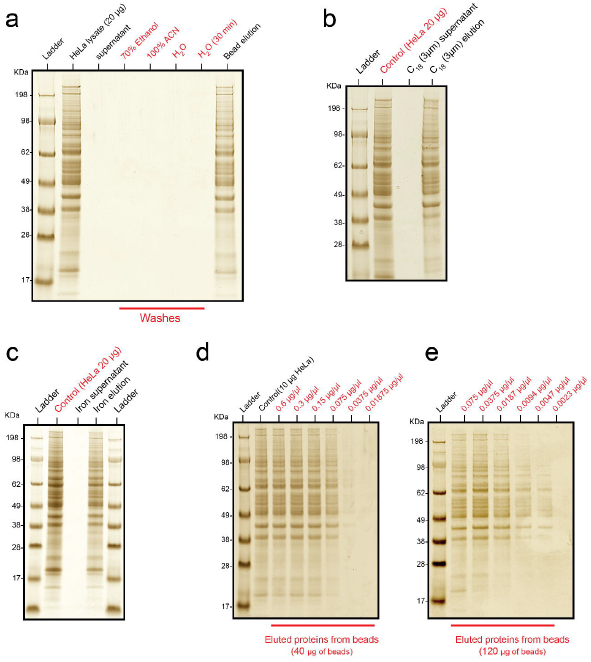
A) Acetonitrile was added to 20μg of HeLa lysate (in 1% SDS) at a final concentration of 70% and the supernatant including sequential washes (including a final 30 minute milli-Q water wash) were analyzed by SDS-PAGE. B) Protein aggregation on non-magnetic microparticles (3μm hydrophobic beads) was analyzed by SDS-PAGE of HeLa lysate aggregated by acetonitrile in a similar fashion. Beads were separated from supernatant by brief (30 second) centrifugation on a benchtop centrifuge. C) Aggregation of 20μg of HeLa lysate (in 1% SDS) on carbonyl iron powder was assessed. After inducing aggregation with acetonitrile, the magnetic powder was separated using a magnet the supernatant was analyzed by SDS-PAGE. Eluate was obtained by adding LDS buffer to the powder and analyzed (after heating at 80°C) by SDSPAGE. D) Recovery of 10μg of HeLa lysate from 40μg of carboxyl coated microparticles were analyzed by SDS-PAGE after the lysate was diluted to decreasing concentration with addition of water and acetonitrile (70% final). E) The amount of microparticles were increased and the capture and elution of highly dilute HeLa lysate from the beads was analyzed by SDS-PAGE.

**Supplementary Figure 2.**
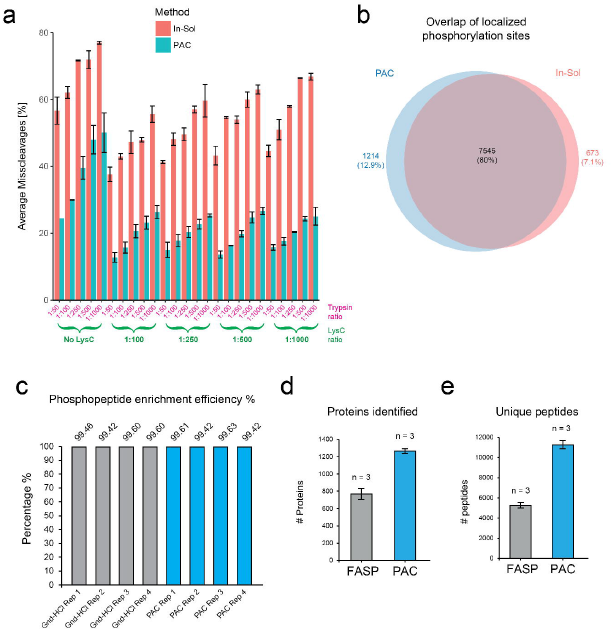
A) Data from Figure 2A is presented in bar plots indicating the percentage of peptides containing missed cleavages at arginine or lysine after digestion with trypsin and lys-c proteases in different combinations and ratios. B) Overlap of localized phosphorylation sites (localization probably ≥0.75) between the two experiments. The site had to be identified in two of the four replicates in both experiment for it to be valid. C) Phosphopeptide enrichment efficiency percentage is shown for all the experiments and replicates in the data presented in Figure 2B. Efficiency was determined by counting all peptides with or without phosphoryl modification (at serine, threonine, tyrosine) in each experiment and after removal of all contaminating and reverse peptides from the evidence.txt output generated by MaxQuant analysis (see Methods). *Error bars represent standard deviation. D) Proteins identified with iBAQ intensities from MaxQuant ProteinGroups.txt output between the two experiments after single shot MS analysis. E) Unique peptides identified between the two experiments and replicates obtained from the evidence.txt MaxQuant output.

**Supplementary Figure 3.**
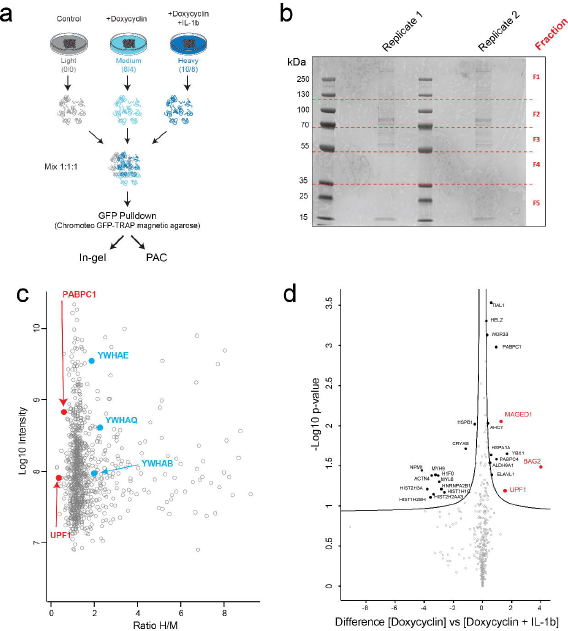
A) SILAC experimental scheme for GFP-TTP pulldown after doxycyclin induction and/or IL-1b stimulation. TPP was expressed with an N-terminal GFP-tag (GFP-TTP) and the bait and interactors were enriched using commercial GFP-Trap beads. Expression TTP with an N-terminal GFP-tag (GFP-TTP) was induced by doxycycline and the interaction dynamics determined after IL-1b stimulation. B) In-gel scheme for generating 5 fractions for in-gel digestion. C) Median of quantile normalized intensities of the proteins were plotted against the average quantile normalized heavy over median (H/M) ratios for all the proteins identified after analysis of GFP-TTP pulldown eluates prepared using PAC. D) Volcano plot of the difference between TTP interacting proteins after 24 hour doxycyclin stimulation (M/L) vs doxycyclin and IL-1b and treated cells at the same time point (H/L). *Error bars represent standard deviation.

**Supplementary Figure 4.**
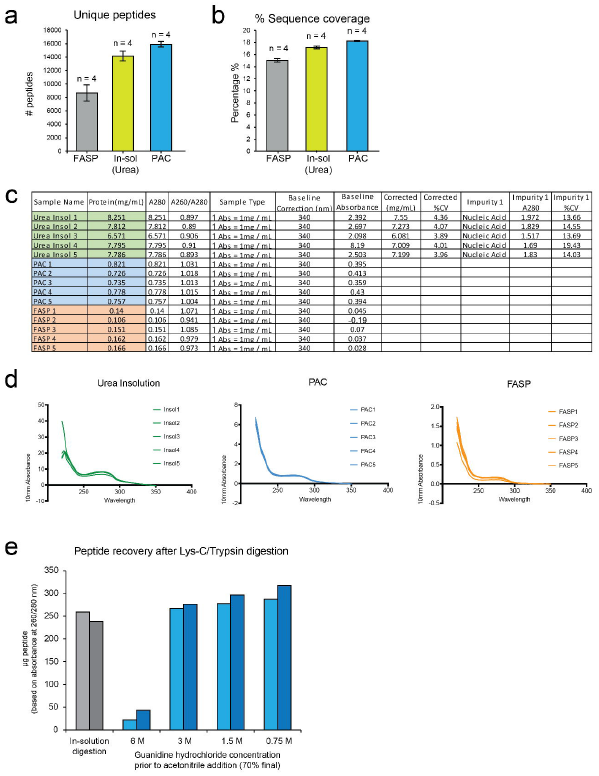
A) Unique peptides identified from the secretome of Raw264.7 cells using different workflows. Unique peptides were counted from the MaxQuant evidence.txt output after removal of duplicate modified sequences as well as reverse and contaminant peptides from each replicate. B) Average sequence coverage was obtained from the identified proteins in each replicate of the secretome workflows. C) Spectroscopy analysis of peptides concentrated and eluted using Stage-Tips after the different secretome workflows shows absorbance and contaminant values at the indicated wavelengths. D) Spectroscopy analysis at different UV wavelengths for the different experiments and replicates is presented. E) Recovery of peptides after lys-c/trypsin digestion using aggregation on microparticles after dilution of the guanidine-hydrochloride lysate with water at different concentrations.

## REFERENCES

1. Wiśniewski, J. R., Zougman, A., Nagaraj, N. & Mann, M. Universal sample preparation method for proteome analysis. Nat. Methods 6, 359 (2009).

2. Shevchenko, A., Tomas, H., Havli\[sbreve], J., Olsen, J. V. & Mann, M. In-gel digestion for mass spectrometric characterization of proteins and proteomes. Nat. Protoc. 1, 2856–2860 (2007).

3. Zougman, A., Selby, P. J. & Banks, R. E. Suspension trapping (STrap) sample preparation method for bottom-up proteomics analysis. Proteomics 14, 1006–1000 (2014).

4. Hughes, C. S. et al. Ultrasensitive proteome analysis using paramagnetic bead technology. Mol. Syst. Biol. 10, 757–757 (2014).

5. Alpert, A. J. Hydrophilic-interaction chromatography for the separation of peptides, nucleic acids and other polar compounds. J. Chromatogr. A 499, 177–196 (1990).

6. Sielaff, M. et al. Evaluation of FASP, SP3, and iST Protocols for Proteomic Sample Preparation in the Low Microgram Range. J. Proteome Res. 16, 4060–4072 (2017).

7. Poulsen, J. W., Madsen, C. T., Young, C., Poulsen, F. M. & Nielsen, M. L. Using Guanidine-Hydrochloride for Fast and Efficient Protein Digestion and Single-step Affinity-purification Mass Spectrometry. J. Proteome Res. 12, 1020–1030 (2013).

8. Ludwig, K. R., Schroll, M. M. & Hummon, A. B. Comparison of In-Solution, FASP, and S Trap Based Digestion Methods for Bottom-Up Proteomic Studies. J. Proteome Res. 17, 2480–2490 (2018).

9. Olsen, J. V. et al. Global, In Vivo, and Site-Specific Phosphorylation Dynamics in Signaling Networks. Cell 127, 635–648 (2006).

10. Wiśniewski, J. R. & Gaugaz, F. Z. Fast and Sensitive Total Protein and Peptide Assays for Proteomic Analysis. Anal. Chem. 87, 4110–4116 (2015).

11. Huttlin, E. L. et al. The BioPlex Network: A Systematic Exploration of the Human Interactome. Cell 162, 425–440 (2015).

12. Ong, S.-E. et al. Stable Isotope Labeling by Amino Acids in Cell Culture, SILAC, as a Simple and Accurate Approach to Expression Proteomics. Mol. Cell. Proteomics 1, 376–386 (2002).

13. Wu, X. & Brewer, G. The regulation of mRNA stability in mammalian cells: 2.0. Gene 500, 10–21 (2012).

14. Chevallet, M., Diemer, H., Van Dorssealer, A., Villiers, C. & Rabilloud, T. Toward a better analysis of secreted proteins: the example of the myeloid cells secretome. Proteomics 7, 1757–1770 (2007).

15. Meissner, F., Scheltema, R. A., Mollenkopf, H.-J. & Mann, M. Direct Proteomic Quantification of the Secretome of Activated Immune Cells. Science 340, 475–478 (2013).

16. Deshmukh, A. S., Cox, J., Jensen, L. J., Meissner, F. & Mann, M. Secretome Analysis of Lipid-Induced Insulin Resistance in Skeletal Muscle Cells by a Combined Experimental and Bioinformatics Workflow. J. Proteome Res. 14, 4885–4895 (2015).

17. Butler, G. S. & Overall, C. M. Proteomic identification of multitasking proteins in unexpected locations complicates drug targeting. Nat. Rev. Drug Discov. 8, 935–948 (2009).

18. Lundby, A. & Olsen, J. V. GeLCMS for In-Depth Protein Characterization and Advanced Analysis of Proteomes. in Methods in Molecular Biology 143–155 (2011).

19. Cox, J. & Mann, M. MaxQuant enables high peptide identification rates, individualized p.p.b.-range mass accuracies and proteome-wide protein quantification. Nat. Biotechnol. 26, 1367–1372 (2008).

20. Tyanova, S. et al. The Perseus computational platform for comprehensive analysis of (prote)omics data. Nat. Methods 13, 731–740 (2016).

